# Biology exams rarely use visual models to engage higher-order cognitive skills

**DOI:** 10.1101/2024.12.23.630136

**Authors:** Crystal Uminski, Christian Cammarota, Brian A. Couch, L. Kate Wright, Dina L. Newman

**Affiliations:** Thomas H. Gosnell School of Life Sciences, Rochester Institute of Technology, Rochester, NY, United States; School of Biological Sciences, University of Nebraska – Lincoln, Lincoln, NE, United States

## Abstract

Visual models are a necessary part of molecular biology education because submicroscopic compounds and processes cannot be directly observed. Accurately interpreting the biological information conveyed by the shapes and symbols in these visual models requires engaging visual literacy skills. For students to develop expertise in molecular biology visual literacy, they need to have structured experiences using and creating visual models, but there is little evidence to gauge how often undergraduate biology students are provided such opportunities. To investigate students’ visual literacy experiences, we surveyed 66 instructors who taught lower division undergraduate biology courses with a focus on molecular biology concepts. We collected self-reported data about the frequency with which the instructors teach with visual models and we analyzed course exams to determine how instructors incorporated visual models into their assessments. We found that most instructors reported teaching with models in their courses, yet only 16% of exam items in the sample contained a visual model. There was not a statistically significant relationship between instructors’ self-reported frequency of teaching with models and extent to which their exams contained models, signaling a potential mismatch between teaching and assessment practices. Although exam items containing models have the potential to elicit higher-order cognitive skills through model-based reasoning, we found that when instructors included visual models in their exams the majority of the items only targeted the lower-order cognitive skills of Bloom’s Taxonomy. Together, our findings highlight that despite the importance of visual models in molecular biology, students may not often have opportunities to demonstrate their understanding of these models on assessments.

## Introduction

The content covered in molecular biology courses exists at a scale so small that biologists rely on visual models, such as drawings and diagrams, to portray the nature of and relationships between invisible molecules. Given the complexity of some of the molecules and molecular processes, biologists often simplify the subject matter by using abstract shapes and symbols to convey meaning in models [1,2]. For example, letters, boxes, and lines can be used to represent concepts such as nucleotides, genes, and chromosomes. Correctly interpreting the abstract shapes and symbols depends on both content knowledge and contextual clues in the representation [3]. Consider that a single line in different contexts could represent the sugar-phosphate backbone of a few nucleotides in a ladder representation of DNA structure, thousands of nucleotides as the line of a box-and-line representation of a gene, or millions of nucleotides in a representation of an unreplicated chromosome. As molecular biologists use abstraction as a form of shorthand, it is critical for students to develop the visual literacy skills necessary to “read” the correct interpretation of the shapes and symbols commonly used to communicate in molecular biology.

Visual literacy is defined as the ability to interpret, create, and communicate with visual models [4], and it is a set of skills that are so important for biology students that they are encapsulated within multiple frameworks for science education. Visual literacy skills are encompassed within the scientific practice “Developing and Using Models” that is part of the three-dimensional framework for science education [5] and are reflected in several core competencies from the *Vision and Change* framework for undergraduate biology education, such as “Ability to use modeling and simulation” and “Ability to communicate and collaborate with other disciplines” [6]. Evidence suggests, however, that communicating science through visual methods is far from standard in biology curriculum [1]. Even though most biology students see visual models in textbooks and lecture slides, visual literacy skills are rarely explicitly taught in biology courses [7].

While it can be difficult to measure the frequency that visual literacy skills are taught without directly observing instructors during class sessions, self-reported data about teaching practices and summative assessments (e.g., tests and exams) can provide insights into how instructors incorporate visual literacy into their courses. The Measurement Instrument for Scientific Teaching (MIST) [8] is one instrument that can be used to gather self-reported data about teaching practices. The MIST specifically includes an item in which instructors can report the frequency that they ask their students to make or interpret models to summarize scientific processes. Models, as broadly defined by the MIST, include drawings, diagrams, schematics, concept maps, flow charts, illustrative models, and box-and-arrow diagrams [8]. Making and interpreting models are key visual literacy skills, thus instructor responses to this item can indicate the frequency in which students encounter instruction about visual literacy in their biology courses. The MIST and other similar measures of teaching practices, such as the Three-Dimensional Learning Observation Protocol [9], can provide information about how often students are asked to make and interpret models, but these measures do not provide context about what types of models students are encountering and how specifically students are being asked to engage with the models.

To get a broader picture of visual literacy practices in introductory-level molecular biology courses, we can analyze the content on instructors’ exams. What is included on exams can indicate the type of content instructors have taught in their courses and can signal what content instructors think was the most important for their students to learn [10,11]. Thus, we would expect to see visual literacy skills, such as interpreting figures and creating diagrams, incorporated into summative assessments if these visual literacy skills are prioritized learning outcomes in a course [7]. As such, if interpreting or creating visual models were prioritized learning objectives in a course, we might expect the instructor to have exam questions which ask students to interpret or create models. Instructor priorities may also be reflected in the point value of exam items [12]. Instructors may signal to students the importance of a concept through the percentage of points certain concepts comprise on an exam, either with higher point value associated with a single item or a large number of points across multiple questions about the same topic. Therefore, if models are emphasized on an exam, we can infer that models were likely to also be an emphasized part of the course content.

In addition to determining how frequently students encounter visual models on their exams, it is also important to consider what students are asked to do with those models. Visual literacy skills span from rote memorization of the parts of a familiar diagram to the creation of new visual representations to communicate a new concept or idea, and these visual literacy skills can be mapped onto levels of Bloom’s Taxonomy [4,7,13]. Bloom’s Taxonomy [14,15] is a widely-used framework for categorizing tasks into ones that can elicit lower-order cognitive skills (remembering, understanding, and applying) and higher-order cognitive skills (analyzing, evaluating, and creating). These higher-order cognitive skills are often equated with critical thinking and tend to be the types of skills instructors report wanting their students to know by the time they finish a course or a degree program [16–19]. By examining the cognitive skills of items with models, we can determine if students are mainly being asked to use models to demonstrate rote content knowledge or if they are being asked to use models to engage higher-order cognitive skills in ways that are more aligned to scientific practices and *Vision and Change* core competencies.

Along with determining the frequency with which instructors are using models on their exams and the cognitive skills of such items, we can also categorize the specific types of models that instructors are using. For a subset of this research, we narrow the focus from visual models in a broad sense to more specifically study only models of DNA. We focus here on DNA because it is a central topic in molecular biology courses and there is an existing framework to categorize visual models of DNA. The DNA Landscape framework [2], is a 3-by-3 matrix that shows variation in DNA representations along axes of scale (nucleotide, gene, and chromosome) and abstraction (literal shape, elements of shape and abstraction, very abstract). To develop molecular biology visual literacy skills, it is important for students to see and interpret representations of DNA across levels of scale and abstraction [1,20,21]; however, an analysis of textbook figures suggested that students may be encountering certain representations in the DNA Landscape matrix much more frequently than others [2]. For example, textbooks were more frequently using abstract “X” shaped representations of chromosomes compared to less-abstract representations of chromosomes as maps or as string-like chromatin [2]. As with textbook figures, we anticipate that certain representations of DNA on biology exams may be more common than others, particularly when considering both the typical topics covered in introductory-level molecular biology courses as well as the ease or the challenge of reproducing certain representations in an exam format.

Visual literacy is a critical skill for molecular biology, but there remains little evidence as to how often these skills are taught or assessed in undergraduate courses [7]. Our work here aims to provide a snapshot of visual literacy in molecular biology courses by addressing the following research questions:

1) How often do students encounter visual models on exams in introductory-level molecular biology courses?
2) What cognitive skills are assessed when items from introductory-level molecular biology exams include a visual model?
3) How do instructors assess student understanding of visual representations of DNA on introductory-level molecular biology exams?

## Methods

### Survey design and administration

This current study expands on the methods and data reported in our previous studies about lower-division biology exams [12,22]. Briefly, we developed a survey for instructors who taught lower-division biology courses (100- and 200-level and their equivalents). We administered the survey through the online platform Qualtrics to collect course artifacts (i.e., a course syllabus, a summative course exam, the exam answer key) as well as relevant information about the biology course and the context in which it was taught. We collected responses from 111 biology instructors at 100 unique undergraduate institutions in the United States. For this research, we focused our analysis on a subset of instructors who taught courses with focus on cell and molecular biology. This subset contained 66 instructors who taught courses in introductory-level cell and/or molecular biology (n = 32), introductory-level general biology (n = 26), genetics (n = 3), microbiology (n = 2), and cell/molecular biology (n = 2). Our sample included broad representation across institution type as defined by Carnegie classifications (S1 Table) and from instructors across career stages (S2 Table).

In our survey, we asked instructors to self-report on a series of factors potentially related to the structure and design of their exams (for full description of the survey design, see our previous publication [22]). One self-report measure in the survey was an abbreviated version of the Measurement Instrument for Scientific Teaching (MIST) [8,23]. For this study, we focus on responses to MIST Item #39 which asked instructors to self-report on the approximate frequency that their students were asked to make or interpret models to summarize scientific processes. To ensure that instructors were interpreting the term “models” similarly, the MIST item defined models as tools to summarize a scientific process that include but were not limited to pathways, diagrams, schematics, concept maps, flow charts, illustrative models, and box-and-arrow diagrams [8]. Likert-scale responses to the MIST item were normalized to a 0-1 scale based on the maximum scale value for the item [8].

This research was classified as exempt from human-subjects review by the University of Nebraska–Lincoln (protocol 21082).

### Coding for item content

Our sample of 66 lower-division cell and molecular biology exams consisted of 2687 items. We used the point values and numbering schemes specified by the instructor to determine the boundaries of individual items. In line with prior recommendations [24], we coded items that shared a common stem and/or used a sub-part numbering scheme (e.g., 3a, 3b, 3c) as a single clustered item. To normalize item point values across courses, we calculated the percentage of total points by dividing individual item point values by the total number of points on the exam and multiplying by 100.

As previously reported for this dataset [12], we coded individual exam item content using existing coding protocols. We used the Three-Dimensional Learning Assessment Protocol (3D-LAP) [24] to code for alignment to each of the scientific practices outlined in the *Framework for K-12 Science Education* [5]. The 3D-LAP uses a set of 2 - 4 criteria statements for each scientific practice, and we previously operationalized these criteria statements into a scale to indicate degrees of alignment to a scientific practice [22]. For this current study, we narrow our analysis to item alignment to the scientific practice “Developing and Using Models,” and provide a schematic of our operationalized coding scheme in Fig 1. For some analyses, we further collapse this coding scheme to categorize items based on whether they contain a visual model. Items that are partially aligned, mostly aligned, or fully aligned to “Developing and Using Models” contain a visual model. Items that are not aligned to any of the “Developing and Using Models” criteria do not contain a visual model.

**Fig 1.**
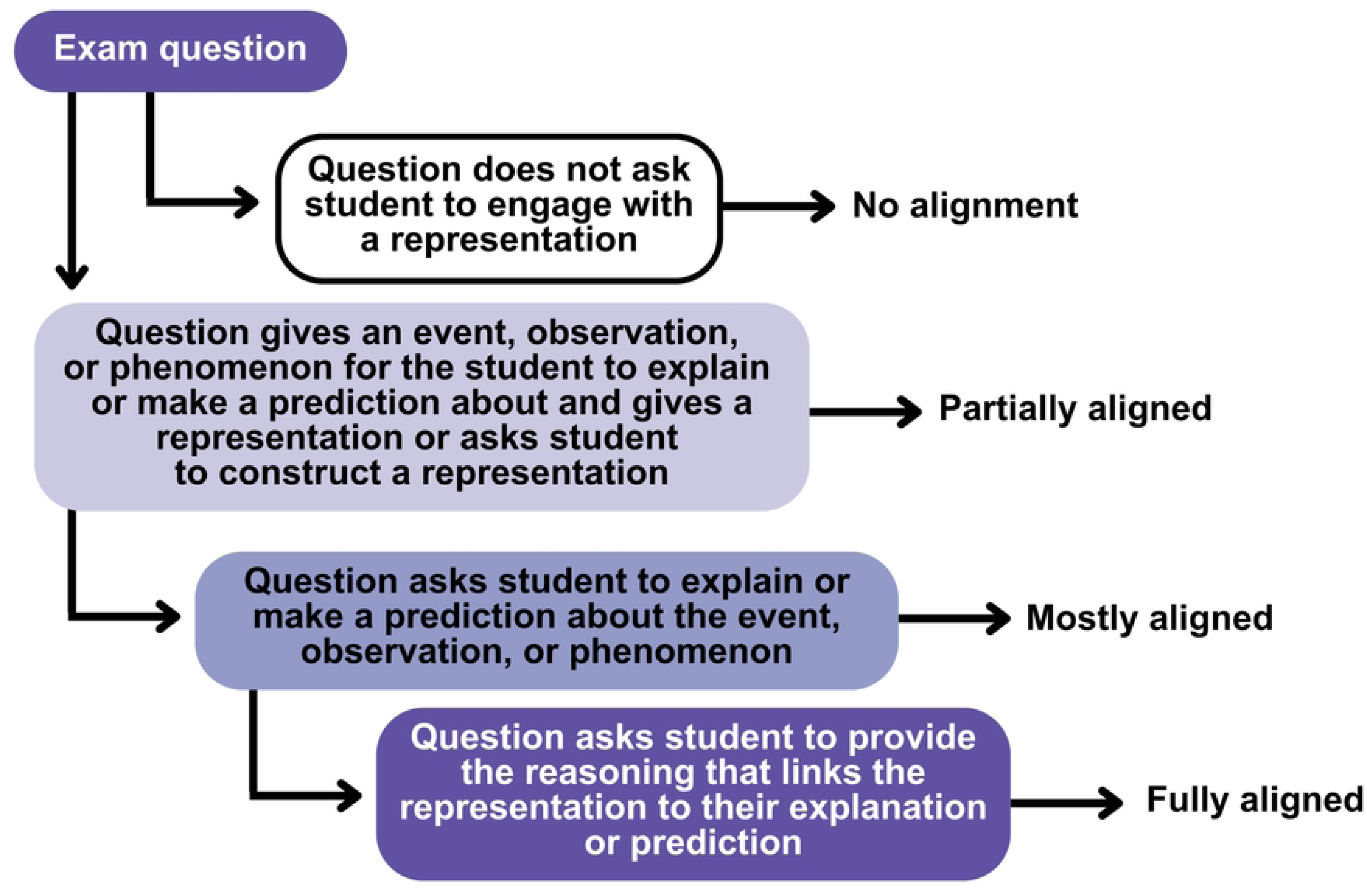
Operationalized 3D-LAP coding scheme for the scientific practice “Developing and Using Models”. We coded each exam item that asked students to engage with a visual model as being “partially aligned,” “mostly aligned,” or “fully aligned” to the 3D-LAP criteria [24] for “Developing and Using Models.” We coded items that did not ask students to engage with a visual model as “No alignment” to “Developing and Using Models.” The 3D-LAP criteria for selected-response and constructed-response items differed in how they describe student interactions with the item (e.g., the criteria indicates that students are asked to construct an explanation rather than select an explanation). The constructed-response criteria statements were operationalized in an identical manner to the selected-response criteria statements here.

The 3D-LAP was designed so that it could be used across science disciplines, and as such, the 3D-LAP criteria for “Developing and Using Models” provides a broad definition of the term “representation” which encompasses mathematical, graphical, computational, symbolic, and pictorial representations [24]. We broadly view the 3D-LAP characterization of “representations” as being equivalent to “visual models,” and note that visual models may be used and applied differently across science disciplines. In the context of biology, we interpreted mathematical representations to include equations when the focus of the model was to identify the symbols or the relationships between variables in the model. If students were provided an equation and asked to use it to perform a calculation, this was coded as a separate scientific practice (“Using Mathematics and Computational Thinking”). We interpreted graphical representations to include models that illustrate generalized trends (e.g., a logistic growth curve). When students were asked to interpret graphs containing actual or simulated data, we coded those items as aligned to a separate scientific practice (“Analyzing and Interpreting Data”). We considered symbolic representations to include letters (e.g., the letters “A,” “T,” “C,” and “G” as symbols for nucleotides), and pictorial representations to include images such as those produced by microscopy. We did not observe any instances of computational models in this sample.

In our prior work [12], we coded items for alignment to levels of Bloom’s Taxonomy [14,15] using the Bloom’s Dichotomous Key [25]. We coded the highest Bloom’s level the item was capable of eliciting and the subsequently categorized the cognitive skills “remember,” “understand,” and “apply” as lower-order cognitive skills (LOCS) and “analyze,” “evaluate,” and “create” as higher-order cognitive skills (HOCS). We treated Bloom’s Taxonomy levels as an ordinal scale for statistical analysis.

Full details on item coding procedures and interrater reliability are in the Materials and Methods of our previous publications on this dataset [12,22].

For this research on the subset of 66 cell and molecular biology exams, we used the DNA Landscape [2] to characterize the visual models of DNA that were present in exam items. Visual models were coded based on the size (nucleotide, gene, chromosome) and the level of abstraction (very abstract, elements of shape and abstraction, literal shape). In some cases, items contained multiple representations of DNA within the same figure (e.g., a gene and a plasmid were both represented in the model). When the same DNA representation was referenced across multiple items (e.g., the figure is used in both exam items 13 and 14), both items were counted as having a representation but the figure is only coded once for alignment to the DNA Landscape. Only representations of DNA were coded for alignment to the DNA Landscape—representations of RNA or microscopy images of DNA were not coded. For a small subset of items, the item stem contextually referenced an image of DNA but the image file was corrupted during data collection efforts. We considered these items to have images of DNA and included them in our statistical analysis, but we did not code these items for alignment to the DNA Landscape.

To establish interrater reliability for alignment to the DNA Landscape, one member of the coding team (CU) first identified all items in the sample that contained a representation of a nucleotide(s), gene(s), and/or chromosome(s). Items that had missing images, contained images duplicated from previous items on the same exam or that contained identical images that were present on multiple exams were removed, leaving a total of 87 unique items in the coding set. Two raters (CU and CC) independently coded the DNA representations in the 87 items for alignment to the DNA Landscape. Coding disagreements were discussed between members of the research team (CU, CC, DLW, and LKW) and consensus values for each item were retained in the final dataset.

### Coding for item format

As described in our previous publication [22], we coded for item format based on response format (constructed-response and selected-response). Briefly, we classified constructed-response items as those that required students to generate an original response and selected-response items as those that asked students to select from a provided set of responses. Constructed-response items included fill-in-the-blank, short-answer, and essay. Selected-response items included multiple-choice, true-false, multiple true-false, and matching.

### Statistical analysis and data availability

We conducted statistical analysis in R (v 4.3.3) using tidyverse [26] for data processing and visualization. We used lme4 [27] to conduct our generalized linear mixed model. As we had multiple items per instructor in the sample, we included instructor as a random effect in our generalized linear mixed model. De-identified data necessary to recreate the generalized linear mixed model and summary statistics presented in the results are in S1 Data.

## Results

### Biology exams rarely ask students to engage with visual models

We analyzed exams from 66 lower-division cell and molecular biology courses and found that exams rarely ask students to engage with visual representations. Of the 2687 items in the sample, only 16% (n = 435 items) provided the opportunity for students to engage with a visual model (Fig 2). While there were very few items with visual models in comparison to the total number of items in the pool, most exams (91%, n = 60) contained at least one visual model.

**Fig 2.**
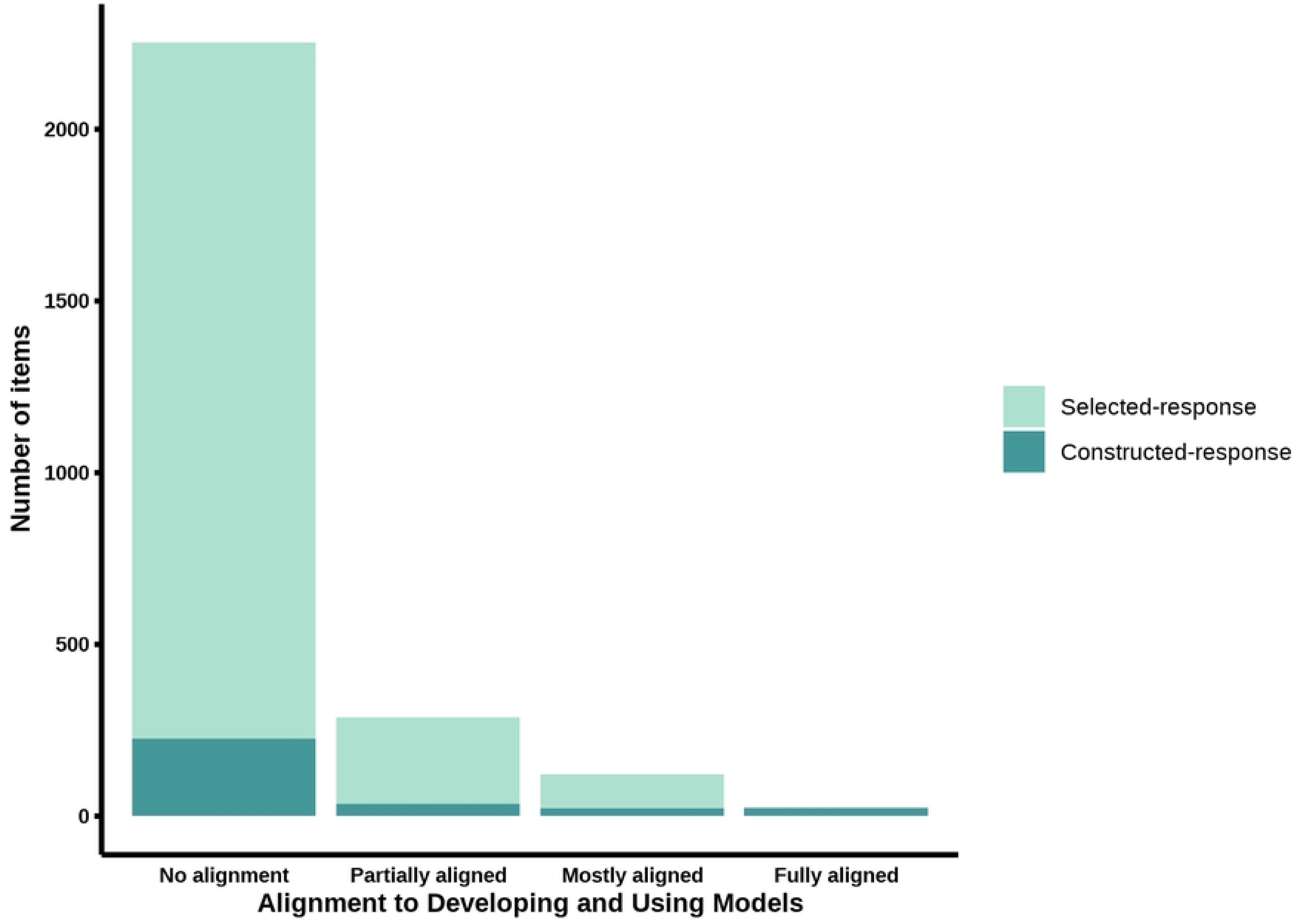
Alignment of cell/molecular exam items to the scientific practice “Developing and Using Models”. Exam items are color coded to distinguish between selected-response and constructed-response format.

Of the 435 items that contained a visual model, the majority of those (66%, n = 288) were only *partially aligned* to the scientific practice “Developing and Using Models.” At this level of partial alignment, items include a representation, but students are only asked to engage with the model at a superficial level, such as labeling parts of a familiar diagram (e.g., identifying organelles in a model of a cell). About a quarter of the items containing a visual model (28%, n = 121) were *mostly aligned* to the scientific practice “Developing and Using Models.” These items asked students to engage with representations to explain a phenomenon beyond memorizable facts but did not explicitly ask students to select or provide the specific reasoning to justify their explanations. Only 6% of items with visual model (n = 26) asked students to explain a phenomenon using appropriate reasoning in ways that *fully aligned* with the scientific practice “Developing and Using Models.”

Items that contained visual models that were *partially*, *mostly*, or *fully aligned* to “Developing and Using Models” were much more often written in a constructed response format than items without models (Fisher’s exact two-tailed test, p < .001). Only 10% of items without models were constructed-response (225 of 2252), yet nearly 90% of items that fully aligned to “Developing and Using Models” were constructed-response items (23 of 26).

When an instructor included a visual model in an item, the items tended to be worth a significantly greater percentage of exam points compared to items without models (β = 0.41, t = 4.4, p < .001; S3 Table). Across instructors, items with models were worth on average 3.0% of exam points (SE = 0.17) compared 2.4% (SE = 0.06) for items without models.

### Self-reports on frequency of teaching with models does not predict use of visual models on exams

On average, instructors assigned 19.6% of exam points (SE = 2.09) to items containing visual models. We compared the percentage of exam points associated with visual models to instructor responses to an item from the Measurement Instrument for Scientific Teaching [8] in which instructors self-reported the frequency with which their students were asked to make or interpret visual models (Fig 3). We found that self-reported frequency of teaching with models is not significantly related to the percentage of exam points with visual representations (Type II ANOVA: F(1,64) = 2.95, p = 0.09).

**Fig 3.**
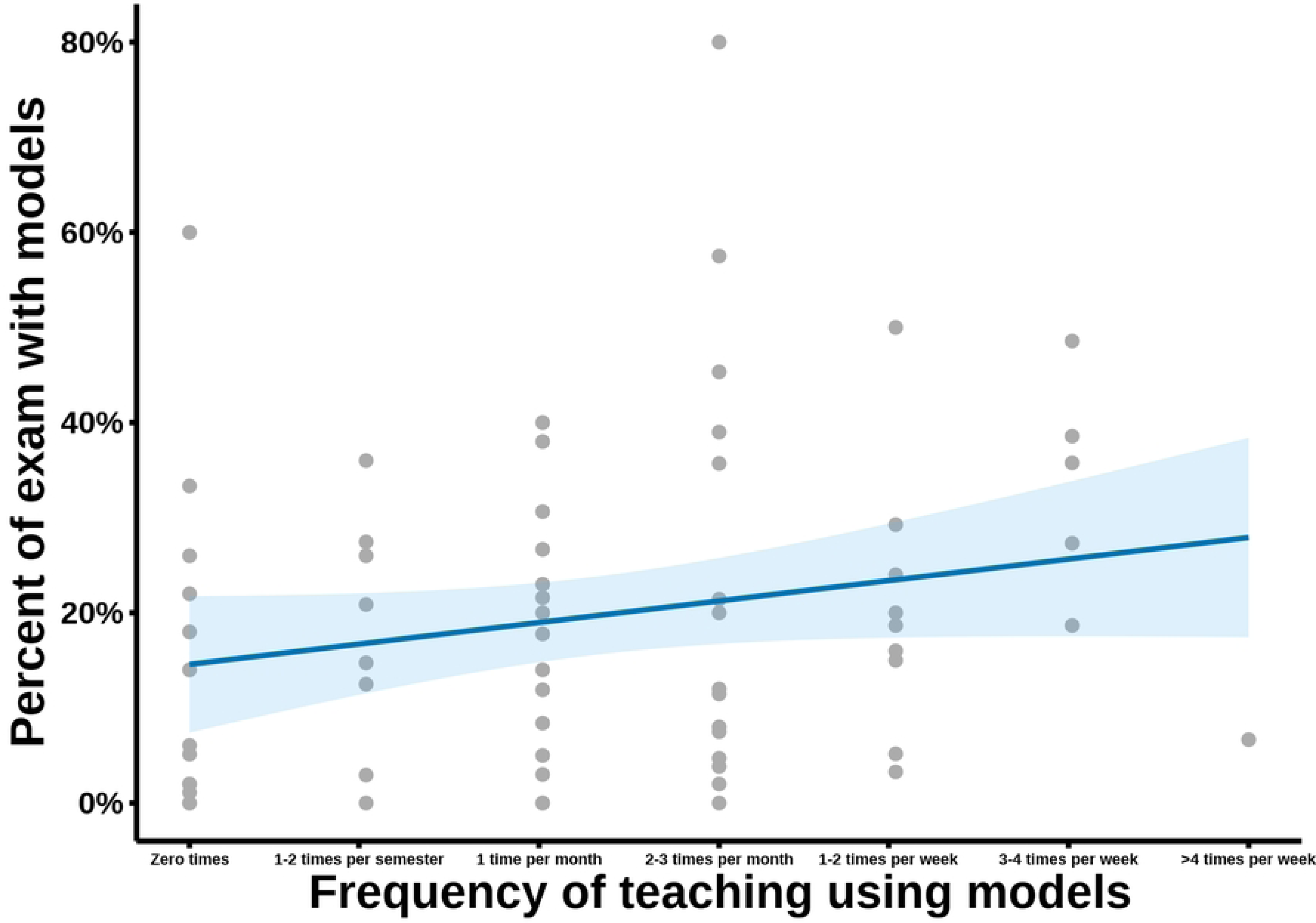
Relationship between self-reported frequency of teaching using models and the percent of exam points that contain a visual model. Teaching practices were self-reported on item #39 from the Measurement Instrument of Scientific Teaching (MIST) [8]. Likert-scale responses to the MIST item, ranging from zero times per semester to more than 4 times per week, were converted into a 0-1 scale, with higher values indicating a greater frequency in which students were asked to make or interpret models during course instruction. Each dot on this plot represents an individual instructor.

### Items with models are slightly more likely to assess higher-order cognitive skills

As we previously described [12], the majority of items in this dataset only targeted the lower-order cognitive skills “remember,” “understand,” and “apply.” Here, we wanted to see if the distribution of lower-order and higher-order cognitive skills differed between items that did and did not have visual models. We compared the proportion of items with and without visual models at each cognitive skill level (Fig 4). The majority of items with models (87%; 378 of 435) and without models (96%; 2157 of 2252) assessed lower-order cognitive skills; however, a chi-square test indicates a statistically significant association between containing a model and testing higher-order cognitive skills, suggesting that these two item features are not independent [ꭓ^2^(1) = 53.9, p < . 001].

**Fig 4.**
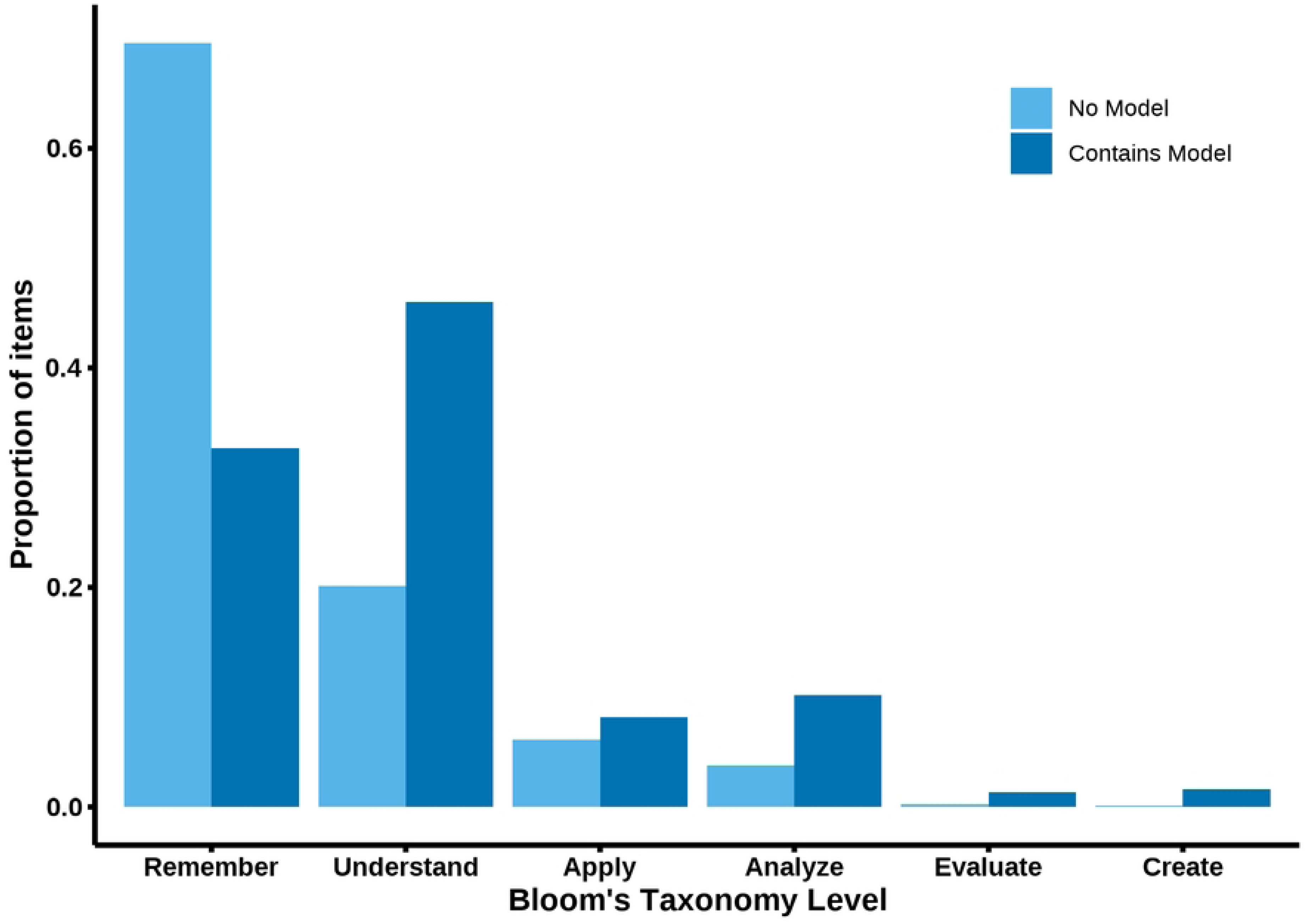
Proportion of cognitive skills assessed by items that did not and did contain visual models. Proportions calculated based on the total number of items with no model (n = 2252) and the total number of items that contained a model (n = 435).

Although there were very few items targeting the higher levels of Bloom’s Taxonomy, these items tended to have a greater degree of alignment to the criteria for the scientific practice “Developing and Using Models” (Fig 5). There is no significant difference between the Bloom’s level of items that were not aligned or were only partially aligned to “Developing and Using Models” (Table 1). In contrast, items mostly aligned or fully aligned to the scientific practice had significantly higher Bloom’s levels compared to items without models.

**Fig 5.**
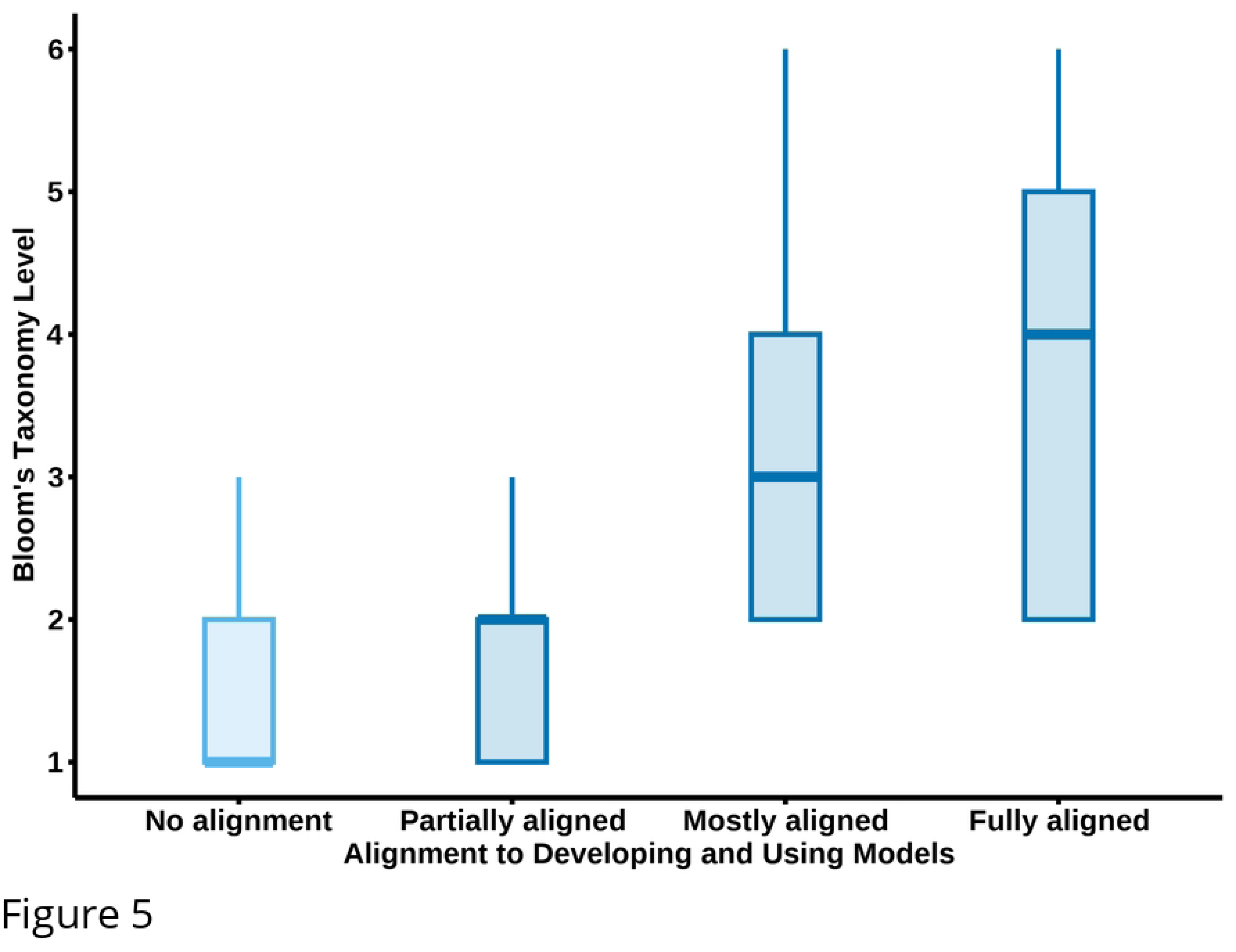
Box-and-whisker plots of the Bloom’s Taxonomy levels of items at each degree of alignment to the scientific practice “Developing and Using Models”. Solid bars within each box represent the median value, boxes represent the interquartile range, and whiskers represent 1.5 times the interquartile range. Bloom’s Taxonomy levels 1–6 correspond to the cognitive skills “remember,” “understand,” “apply,” “analyze,” “evaluate,” and “create,” respectively.

**Table 1.** Linear regression model predicting the Bloom’s Taxonomy level of items based on alignment to “Developing and Using Models”.

| Fixed effects | $\beta$ | Standard Error | $df$ | $t$ | $p$ |
| --- | --- | --- | --- | --- | --- |
| Intercept | 1.57 | 0.06 | 44 | 26.3 | < .0001 |
| Partial aligned | 0.09 | 0.05 | 2638 | 1.9 | 0.052 |
| Mostly aligned | 1.10 | 0.07 | 2625 | 15.5 | < .0001 |
| Fully aligned | 2.04 | 0.15 | 2667 | 13.2 | < .0001 |
| Model: Bloom's level ~ Alignment to "Developing and Using Models" + (1\|instructor)<br>Reference group: No alignment<br>$R^2 = 0.36$ | | | | | |

### Most representations of DNA on biology exams are very abstract

We examined a subset of exam items with visual models of DNA. Across the 66 cell and molecular biology exams in the sample, 55% of exams (n = 36) had at least one representation of DNA. In total, there were 146 exam items with DNA representations which comprised only 5% of the total number of items in the sample. Of these, 87 contained a unique representation and 46 duplicated images from prior items on the same exam. There were an additional 13 items from exams administered through an online learning management system (e.g., Canvas, Blackboard) for which the item stem indicated an image but the image files were corrupted during data collection efforts.

We categorized the 87 items with unique representations of DNA for their alignment to the DNA Landscape. Most of the representations of DNA aligned to a single location on the DNA Landscape, but there were 12 items that contained two representations, and as such there were a total of 98 DNA representations in the sample. We found that there were examples of DNA representations at each scale of nucleotide, genes, and chromosomes, but across these scales, most of the representations were very abstract (Fig 6). The majority of the images (59%, 58 of 98) were very abstract representations in which nucleotides, genes, and chromosomes were represented with letters and symbols. Notably, there were not any representations of gene maps in which the representation included the location of an actual gene or the distance between multiple genes on a chromosome.

**Fig 6.**
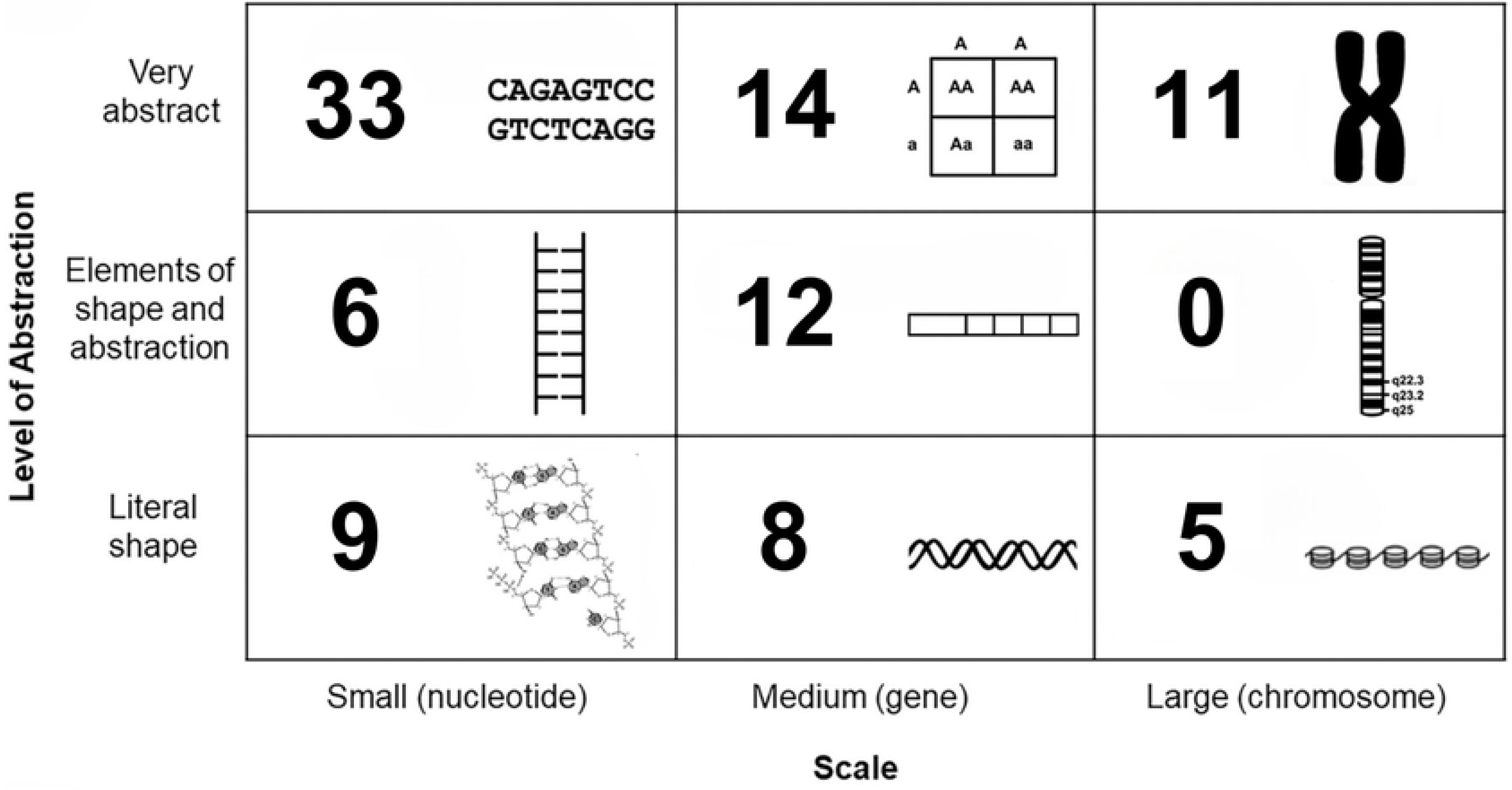
Categorization of the representations of DNA in molecular biology exams based on their alignment to the scale and abstraction axes of the DNA Landscape. There were unique DNA representations on 87 exam items, and 12 of those items had representations that were coded in multiple places on the DNA Landscape.

The majority of items with representations of DNA tended to assess lower-order cognitive skills (93%, n = 136). In comparison, while the majority of items with non-DNA representations also assessed lower-order cognitive skills (83%, n = 239), there was a statistically significant difference between the proportion of lower-order and higher-order skills assessed between the items with DNA and non-DNA representations (Fisher’s exact two-tailed test, p = .004). Items with representations of DNA often asked students to complete rote tasks, such as selecting a sequence of complementary nucleotide base pairs, matching representations of mitotic stages, or identifying nucleotides within a series of monomer chemical structures. When comparing the cognitive skills assessed, the items with representations of DNA were statistically different from the rest of the larger pool of items with representations.

## Discussion

Using models is a key scientific practice and competency for biology students [5,28] and visual literacy skills appear in nearly a third of nationally-endorsed learning objectives for introductory biology [29], yet we found that these skills are rarely assessed on undergraduate biology exams. Thus, our findings underscore the disconnect between the goals and priorities of national calls and what is practiced in undergraduate biology courses. While most introductory-level biology exams predominately assess lower-order cognitive skills, exam items containing models were slightly more likely to ask students to engage in higher-order thinking. Here, we highlight the utility of models as a means to engage students’ higher-order cognitive skills, call for more robust assessment of students’ molecular biology visual literacy skills, and discuss ways to address challenges to using models to teach and assess visual literacy.

### What might be constraining assessment of visual literacy skills in introductory biology?

Our analysis revealed that introductory-level biology exams rarely asked students to use visual literacy skills to engage with models. Overall, there were very few items that included a visual model. Even though our findings indicate that visual models can be a means to engage higher-order cognitive skills, many items containing a model only targeted memorization and recall. We found that when instructors included visual models on their exams, many of those items were only asking students to label familiar structures. While labeling can help students learn about molecular structures [30], labeling alone does not address the suite of visual literacy skills that is needed for expertise in molecular biology. To gauge students’ visual literacy and ability to use higher-order thinking, instructors should create assessments that ask students to evaluate the power and limitations of models, construct models to explain a concept, translate between multiple representations of the same concept, or to compare representations at various scales [21]. For example, an instructor could present students with an item that contains a written description of a particular point mutation and ask them to explain why an abstract visual model of a chromosome is not an effective model to represent the mutation and to draw a new model that would better represent the mutation. Such an assessment item would ask students to evaluate the limitations of a chromosome model, construct a more appropriate model to explain a mutation, ask students to translate between written and graphical representations of a mutation, and compare models of DNA across different scales. Despite the importance of these visual literacy skills in biology, we found that very few exams specifically targeted these skills.

One potential explanation for these results is that instructors have very few examples of what a visual literacy assessment “looks like.” While the Drawing-to-Learn framework provides a robust set of recommendations for facilitating assessment of visual literacy skills [7], there are few examples of visual literacy assessments that instructors can readily use or adapt in their biology courses. Instructors who rely on assessments generated by curriculum publishers may encounter the challenge that most figures in “test bank” items are often only asking students to add labels to a diagram. Instructors may turn to published biology concept assessments or concept inventories (e.g., [31–35]), but models are rarely the focus of such assessments and students may be interpreting the models on concept assessments in different ways than the assessment developers intend [36]. To date, there are no valid and reliable assessment instruments that are specifically intended to measure a suite of visual literacy skills in molecular biology. We emphasize the need for biology education researchers and assessment developers to create assessment instruments that can be used to measure biology visual literacy skills. Such visual literacy assessment instruments may be directly used by biology instructors in their classes or may serve as inspiration for how instructors can create their own visual literacy assessments.

We also consider our results in light of the limitations that often constrain how instructors are able to design and administer their exams. Instructors often have limited time to write and grade exams [37], and it can take a substantial amount of time to write and grade three-dimensional exam items that address scientific practices (such as the practice “Developing and Using Models”) [38]. Grading exams can be especially time consuming when instructors use constructed-response (i.e., open-ended) items where students can create models and articulate their reasoning in a written format. When asking students to create their own models on exams is not feasible, instructors may assess student understanding of models if they integrate pre-fabricated models as multiple-choice options.

Another potential explanation of our findings is that instructors may be limited in their ability to locate, create, or reproduce models for an exam context. To ensure that students are engaging visual literacy skills and not simply memorizing a familiar diagram for an exam, instructors may need to locate figures that are similar but different from the exact model they have used in class, which can be challenging. There may be material and financial constraints involved with creating physical copies of the figures for in-person exams, especially if the figures include color. While there are certainly time and resource constraints in grading constructed-response questions, having students create their own models to demonstrate their understanding of visual literacy conventions circumvents the need for instructors to invest time and resources into locating, creating, or reproducing models themselves.

### Backward Designing biology curriculum to emphasize visual literacy in both teaching and assessment

We found that there was a mismatch between the frequency with which instructors reported teaching using models and the percent of their exams that contained models. In some cases, instructors who reported that they frequently taught using models (≥ 1-2 times per week) rarely incorporated models into their exams, while conversely, some instructors who reported never directly teaching models had used models on upwards of a third of their exam. This mismatch may reflect how instructors are thinking differently about teaching and assessing using models, and we anticipate that this mismatch can lead to confusion and frustration for students. If exams are misaligned to instruction, students may be left wondering why they spent time on content they were not tested on and they may question if it is fair that they were tested on content they were never directly taught. We recommend taking a Backward Design approach to curriculum development to address this mismatch between instruction and assessment of modeling. Following principles of Backward Design, course learning goals should be reflected in what is assessed and what is taught in a course [39]. Accordingly, if using models and visual literacy skills are prioritized learning outcomes in a biology course, modeling and visual literacy should be incorporated into exams and integrated into instruction. Instructors may also use point value of exam items to signal to students that modeling and visual literacy are priorities. Higher point values on items with models may help to strengthen students’ association that modeling is a valuable and important skill for them to learn.

### Using models is a method to engage students in higher-order thinking

Even though visual models have the potential to be powerful tools to engage student reasoning with higher-order cognitive skills, when instructors incorporated visual models into their exam items, these items were often only targeting lower-order cognitive skills, such as labeling structures on a familiar diagram. While items with models can certainly be used to assess higher-order cognitive skills, our findings highlight that instructors may not be capitalizing upon using exams to engage students in model-based reasoning. We encourage instructors to consider ways to ask their students to critically analyze and evaluate the models they are presented. For example, instructors can ask students to explain how and why a biologist might use abstraction to simplify a model and why such simplifications may be useful for conveying certain information. Instructors can also ask students to consider when such simplifications obscure some relevant information and when alternate models may be more useful in a particular context.

We also encourage instructors to consult the 3D-LAP criteria statements for “Developing and Using Models” (reproduced in Fig 1) as methods for designing exam items that target critical thinking. A key feature of the 3D-LAP criteria is the emphasis on asking students to productively use models to engage in reasoning. Following the 3D-LAP guidelines, it is not only important for students to be able to use a model to explain a phenomena—students need to be able to explain the reason *why* that model is useful for explaining a phenomenon. This reasoning component provides important information for instructors to gauge how their students are thinking about models, yet this reasoning component is often absent from exam items [22,40,41]. As full alignment to the 3D-LAP criteria for “Developing and Using Models” was significantly associated with higher levels of Bloom’s Taxonomy, we suggest instructors consider ways to ask students to demonstrate their reasoning. Instructors who have the time and resources to grade open-ended questions can incorporate explicit directions in their items to elicit reasoning in a written format. Students may be more likely to demonstrate reasoning about models if they are prompted to describe how they arrived at a conclusion or to support their argument using evidence. If instructors are writing closed-ended questions, we recommend formats that enable multiple points of student input, such as multiple-true-false questions [32,42,43] or two-tiered questions that allow students to first use a model to explain a phenomenon and then complete a follow-up question in which they can demonstrate their reasoning.

### Emphasizing the value of abstract models in biology

We were not surprised to see that most representations of DNA in our sample were abstract models. Biologists frequently use abstract models because the shapes and symbols in abstract models highlight the most salient structures and features in simple-to-draw ways. These abstract models are often easy for instructors to recreate for an exam context and tend to be well-suited for grayscale printing. Instructors can use abstract models to create meaningful opportunities for model-based reasoning [7], yet we saw few examples of this in how abstract models were used on exams. Exam items with abstract representations mostly focused on rote tasks, such as matching and labeling. In both teaching and assessment, we encourage instructors to consider how to leverage the value of abstract models and ask students to think deeply about the purpose and efficacy of such representations. For example, in addition to asking students to use letters to complete a complementary strand of nucleotides, instructors can ask students to explain the reason why letters are a useful model for that context (as compared to the ladder model or the nucleotide chemical structures). Instructors can alternatively ask students to explain a situation when letters are not a particularly useful model. Students often encounter abstract models, so we encourage instructors to consider ways to ask students to engage with *why* and *how* those abstract models convey meaning in biology.

### Limitations

We relied on exams to make inferences about what content students were encountering in their coursework. This methodology is common in education research studies [44–47], but we acknowledge that exams do not capture the full range of assessment types that can be administered in introductory-level biology courses. It is possible that students engaged with models in formative assessments or other types of summative assessments, such as projects or presentations, which were not reflected in our sampling.

## Conclusion

Using models is a key skill for biologists, yet our research identified that introductory-level students often have few opportunities in their course exams to demonstrate this ability. Despite the importance of visual models in molecular biology, models are rarely incorporated into introductory-level molecular biology exams. When models are present in exam items, they can have the potential to elicit higher-order cognitive skills yet are often only used to assess students’ ability to recall rote information.

This finding signals a greater need for assessments to incorporate visual literacy skills and to engage students in demonstrating their understanding of why and how models are useful tools in biology.

## Acknowledgments

We thank the instructors who participated in this research. We thank Noah Courtney for contributions in early conceptualization of this project.

## Supporting Information

**S1 Data. De-identified data.**

**S1 Table. Institutional Carnegie classifications.**

**S2 Table. Self-reported teaching experience of undergraduate biology instructors.**

**S3 Table. Linear mixed model predicting the point value of items with visual models.**

